# Gap junction internalization and processing in vivo: a 3D immuno-electron microscopy study

**DOI:** 10.1101/2020.06.29.178475

**Authors:** Rachael P. Norris, Mark Terasaki

## Abstract

Gap junctions have well-established roles in cell-cell communication by way of forming permeable intercellular channels. Less is understood about their internalization, which forms double membrane vesicles containing cytosol and membranes from another cell, called connexosomes or annular gap junctions. Here, we systematically studied the fate of connexosomes in intact ovarian follicles. High pressure frozen, serial sectioned tissue was immunogold labeled for Connexin 43. Within a volume of electron micrographs, every labeled structure was categorized and counted. Surface area measurements indicate that large connexosomes undergo fission. Subsequent modifications are separation of inner and outer membranes, loss of Cx43 from the outer membrane, and outward budding of the modified membranes. We also documented several clear examples of organelle transfer from one cell to another by gap junction internalization. We discuss how connexosome formation and processing may be a novel means for gap junctions to mediate cell-cell communication.

## Introduction

Gap junctions are arrays of permeable channels between two cells that have well-established roles in intercellular signaling (Nielsen et al., 2012). The basic structural unit is the transmembrane protein connexin. Six connexins assemble to form the connexon, which is a pore within the membrane. A connexon in one cell docks head-on with a connexon in a neighboring cell to form a channel between the two cells. Large gap junctions may consist of hundreds or thousands of channels packed densely in a patch a few microns in diameter (Larsen, 1977).

The current view is that connexons are added to the plasma membrane via small post-Golgi vesicles, followed by docking between two cells, but how connexons are removed from the membrane and their subsequent fate are incompletely understood. To turn over the gap junction, cells could undock connexons and then endocytose them in small parcels. Instead, connexons remain docked and the gap junction is taken up by one of the two cells (Laird et al., 2006; Falk et al., 2016). This was first suggested by electron microscopists who interpreted circular gap junction profiles (first called annular gap junctions) as internalized gap junctions (Espey and Stutts, 1972; Merk et al., 1973). This interpretation was convincingly corroborated by live imaging of GFP-connexins, which showed formation of vesicles of comparable size (Jordan et al., 2001; Piehl et al., 2007). The internalized gap junction structure is now often called a connexosome (Laird, 2006).

Gap junction internalization is therefore a type of endocytosis, in which the plasma membrane of the neighboring cell remains attached to the endocytosed plasma membrane (Heck and Devenport, 2017). Gap junction internalization can also be considered to be a form of trogocytosis (Joly and Hudriser, 2003) in which a portion of the plasma membrane and cytosol of the neighboring cell is transferred to the engulfing cell (See Fig. 1A). There is a possibility that further processing after the initial engulfment is involved in other modes of cell-cell communication (see Discussion). Internalized gap junctions could likewise play new roles in cell-cell communication.

**Figure 1.**
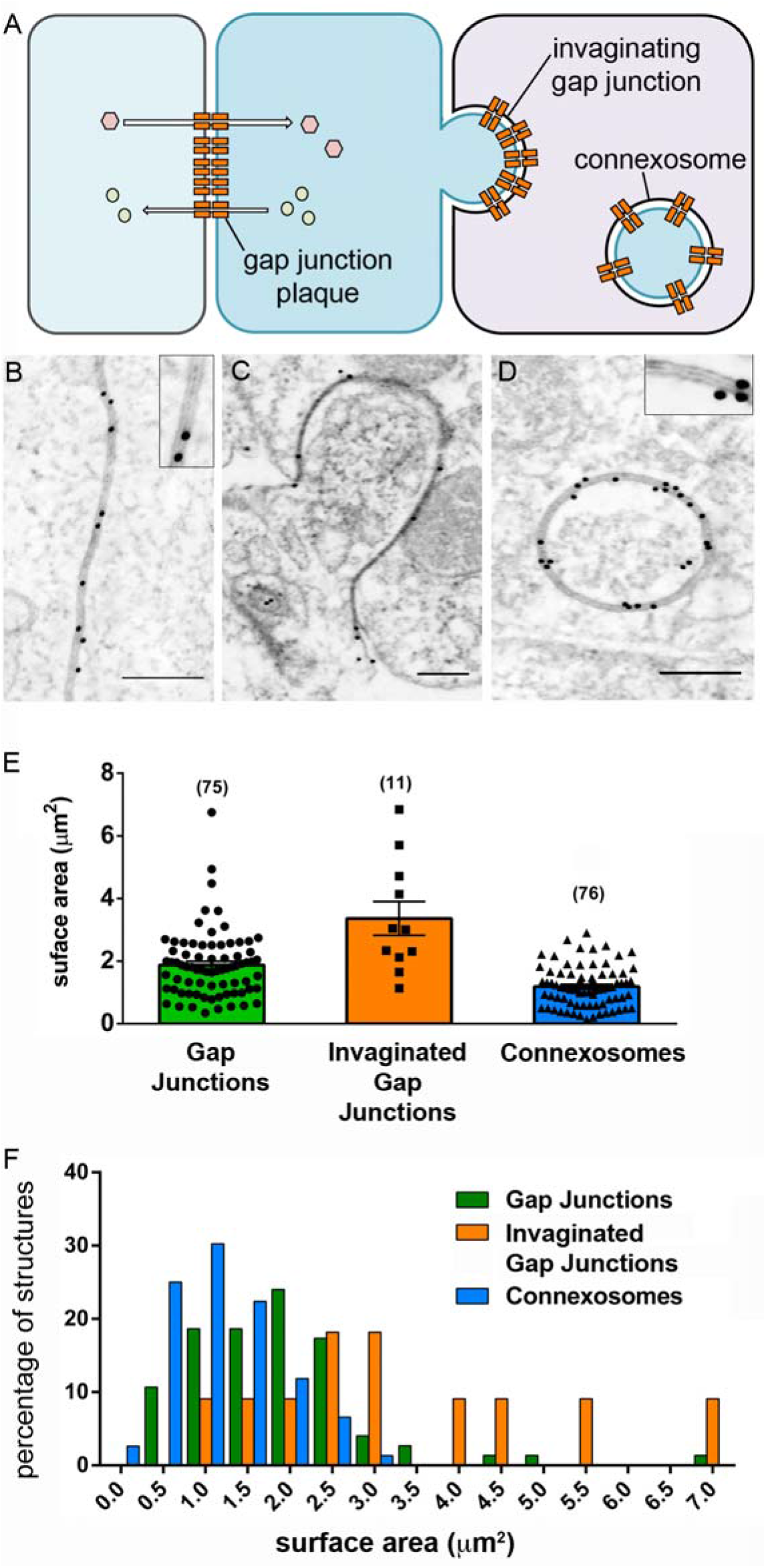
Immunogold labeling and surface area measurements of different forms of gap junctions. **A.** A schematic of a gap junction plaque, an invaginating gap junction, and a fully internalized gap junction (connexosome). Gap junction proteins (connexins) are represented by orange rectangles. The connexosome contains cytoplasm and membrane from the neighboring cell, depicted in blue. **B-D.** Electron micrographs of a gap junction plaque (B), an invaginated gap junction (C), and a connexosome (D). Gap junctions are labeled with an anti-Cx43 antibody and 10 nm-gold conjugated secondary antibody. The multi-laminar structure of gap junctions is visible at high resolution (insets). Scale bars are 250 nm. **E.** Surface areas of all gap junction plaques, invaginated gap junctions, and connexosomes that were measured in a volume of ~30,000 μm^3^. **F.** Surface area measurements plotted as a histogram. The binning interval was 0.5 μm^2^. For each bin, the percentage of the total number of structures is graphed. The smaller area of connexosomes is consistent with budding or fission of internalized gap junctions.

In a previous study (Norris et al., 2017), we addressed several methodological issues that have limited electron microscopic studies of gap junctions in the past. Mouse ovarian follicles were high pressure frozen in order to preserve structure better than chemical fixation, which involves diffusion of aldehydes through the cell membrane(s) and cross linking of proteins, during which abnormal processes may occur (Murk et al., 2003a). The frozen tissue was freeze-substituted, embedded in lowicryl, sectioned, then immunolabeled with an antibody to Cx43. This allowed us to unambiguously identify gap junctions. Serial sections were then imaged to obtain three-dimensional information; this distinguished between internalized connexosomes and gap junctions in the process of invagination. Likewise, a round profile in a single section could be a vesicle or an invagination, or an apparently empty vesicle could contain intraluminal vesicles; serial sections distinguished these alternatives.

Here, we investigate the fate of internalized gap junctions by examining Cx43 localization. As before, we high pressure froze ovarian follicles and immunolabeled serial sections. We found a surprisingly large number of Cx43 labeled structures that we interpret as modifications of the connexosome.

## Results

Mouse ovarian antral follicles were high pressure frozen 30 minutes after exposure to luteinizing hormone (LH). This tissue was used because it has large numbers of Cx43 gap junctions (Okuma et al, 1996; Norris et al., 2008, 2017; Baena et al., 2020), which are caused to internalize in response to hormone (Larsen et al., 1987). As in our previous study, the follicles were embedded in lowicryl, serial sectioned, then labeled with primary antibody to Cx43 and gold-labeled secondary antibody (Norris et al., 2017). We used a different scanning electron microscope in order to obtain higher resolution images from the sections labeled in our previous study. This provided clearer images of the classic double membrane with a gap structure (Figure 1B-D).

We will use the term connexosome to mean a double membrane vesicle completely detached from the plasma membrane in which both limiting membranes were contacting each other and labeled with Cx43 throughout the periphery of the vesicle. Connexosomes were the most abundant Cx43 labeled structure in the cytoplasm, but there were also many other membranous structures that contained Cx43 (Video 1) which we interpreted to be modifications of the connexosome.

We characterized these Cx43 structures by classifying and counting them in a defined volume. By doing so, we gained information on the relative abundance of various forms, which could be useful in deducing dynamics. We analyzed two volumes of mural granulosa cells that were each 60 μm x 60 μm in the x/y plane (imaged at 5 nm per pixel), in 73 sections of 60 nm thickness. Thus each volume was 15,768 μm^3^. This volume corresponds to approximately 35 spherical cells with a 12 μm diameter.

In this volume, there were 86 gap junctions. Of these, 75 were a planar patch. The 11 others were present on highly infolded membranes, which we refer to as invaginating gap junctions. There were 132 internalized structures. Of the 132 Cx43 labeled structures, 76 were connexosomes and 56 appeared to be connexosomes that had undergone processing.

### Connexosome formation

Direct imaging of living cultured cells has shown that connexosomes are formed by internalization of entire gap junctions, and that connexosomes can undergo fission (Piehl et al., 2007; Bell et al., 2018). We examined the high pressure frozen and serial sectioned ovarian follicles for connexosome formation. The areas of structures are measureable in 3D data sets, so we also measured the surface areas of all of the Connexin 43 containing membranes because this information could be relevant to connexosome formation (Figure 1E).

The average surface area of the 75 “flat” gap junction plaques in the volume was 1.9 ± 1.1 μm^2^ (mean ± SD), with a range of 0.36-6.8 μm^2^. Serial sections showed that most were disc shaped, corresponding to a disc of average diameter 1.56 μm. The 11 invaginated gap junctions looked like they could form connexosomes. These were larger than the “flat” gap junctions, with an average surface area of 3.4 ± 1.8 μm^2^ with a range of 1.1-6.8 μm^2^ (N = 11).

Connexosomes, on the other hand, had an average surface area of 1.20 ± 0.7 μm^2^, and a range of 0.14-6.8 μm^2^ (N=76). The average connexosome area thus was 64% of the average gap junction area (Figs. 1E, F).

It seems likely that invaginated gap junctions were frozen in the process of internalization and are therefore the source of connexosomes. The surface areas of invaginated gap junctions are significantly larger than connexosomes. In live imaging of GFP-Cx43 expressing cells, connexosomes were observed to bud off small vesicles (Piehl et al., 2007; Bell et al., 2018); in the ovarian follicle, the connexosomes may undergo fission.

### Cytoplasm and organelle transfer

Four of the 11 invaginated gap junctions contained organelles or vesicles in addition to cytosol. One deeply invaginated gap junction contained a multivesicular endosome and a mitochondrion (Fig. 2A, Video 2). If such structures were to form connexosomes, it would result in transfer of organelles from one cell to another.

**Figure 2.**
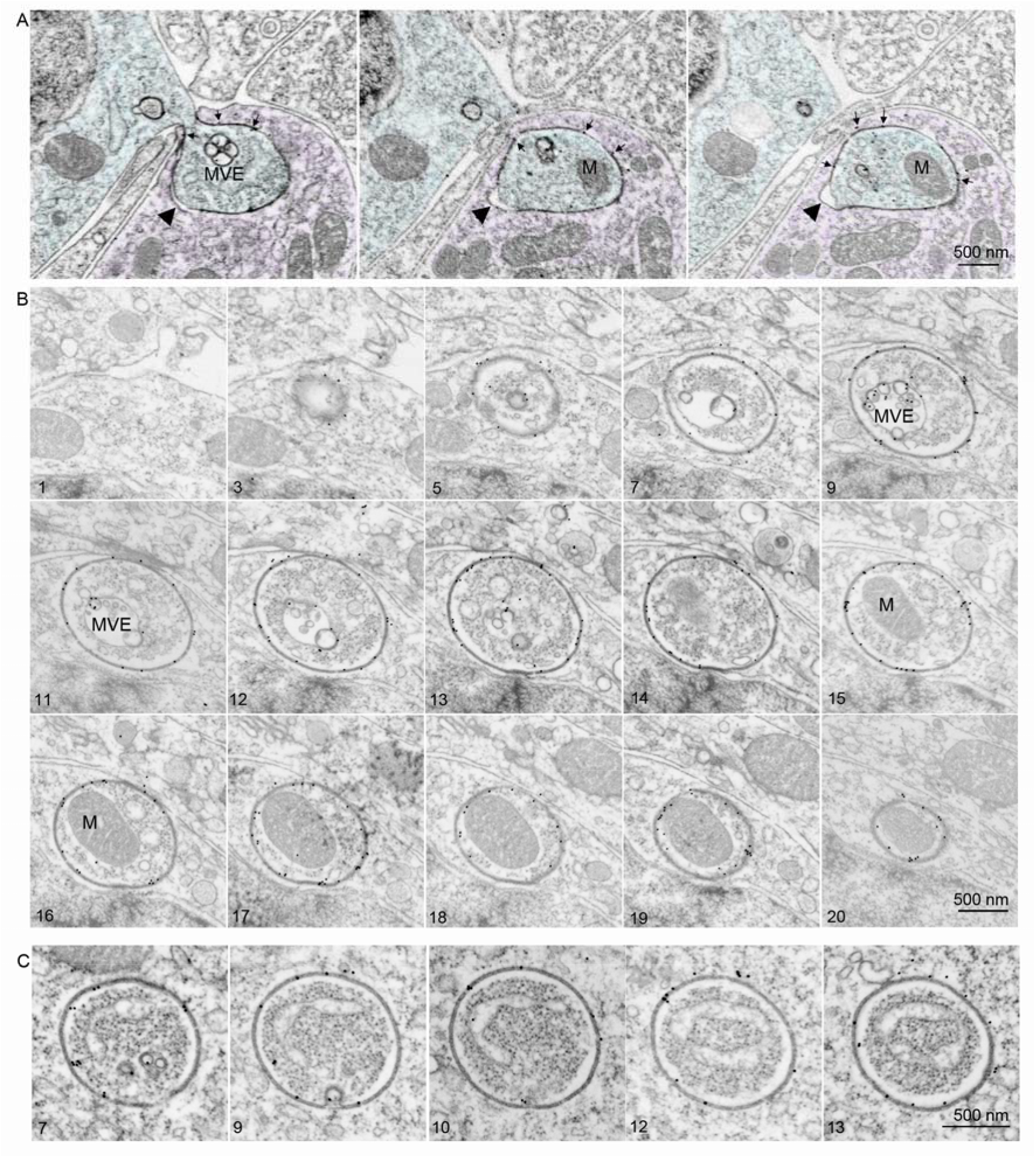
Transfer of cytosol and organelles by way of gap junction internalization. **A**. Three serial sections through an invaginated gap junction containing a multivesicular endosome (MVE) and a mitochondrion (M). Gold particles depicting Cx43 are indicated with arrows. For clarity, the cytoplasm of the cell protruding into another is shaded in blue. The arrowhead indicates a small region of apposed plasma membranes that were not joined together by connexons but were presumably incorporated into a connexosome. **B**. Fifteen serial sections through a connexosome containing a multivesicular endosome (MVE), a mitochondrion (M) and smaller vesicles. The complete series has 22 sections, and the section number is shown in the lower left corner. **C**. Five serial sections through a connexosome containing a tubular organelle. The complete series has 18 sections, and the section number is shown in the lower left corner. Related Video 2 shows the full structures of those depicted in A-C.

We indeed found a connexosome containing a mitochondrion and an apparent endosome (Fig. 2B, Video 2), and another connexosome enclosing a tubular organelle, possibly ER (Fig. 2C, Video 2). Because the entire periphery of the connexosome is gap junction, these organelles must have come from other, previously gap junction-coupled cells. In total, 17 out of 76 connexosomes contained vesicles or organelles in addition to cytosol.

### Connexosome modifications

Of the 132 Cx43 labeled internalized structures, 56 appeared to be modified connexosomes. To describe the modifications, we will use the following terms to refer to the membranes and compartments of the unmodified connexosomes. There are an outer and an inner membrane, which are closely apposed because the connexons are docked. The small space between the two membranes originally was the extracellular space. The compartment within the inner membrane came from the cytoplasm of the neighboring cell.

The initial modification appears to be fusion of a connexosome with another vesicle. In 7 of the 56 modified connexosomes, a patch of unlabeled outer membrane bulged outward from a labeled inner membrane (Fig. 3A, Video 3). This is what would be expected if a vesicle had fused with the outer membrane (Fig. 3B, Video 3). The space between the two membranes was either dark (Fig. 3B) or clear (Fig. 3C) suggesting that different types of vesicles were involved. Cx43 labels the inner membrane in areas where the inner and outer membranes have separated. In these regions, the connexons must have become undocked.

**Figure 3.**
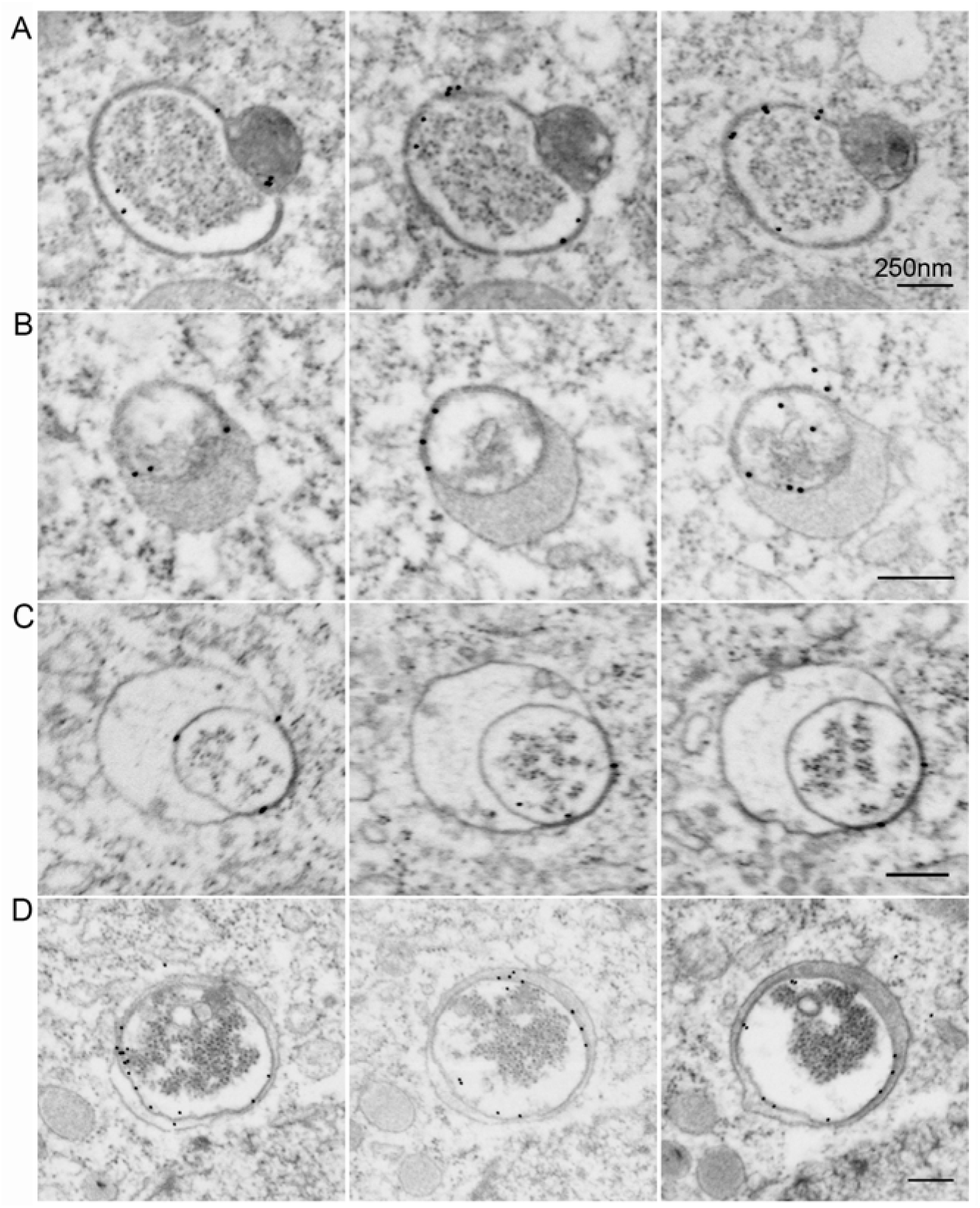
Modifications of connexosome membranes and contents. **A.** Three serial sections showing apparent fusion of a connexosome and a dense vesicle. **B.** Three serial sections through a modified connexosome with an electron dense space between the separating membranes. **C.** Three serial sections through a modified connexosome with an electron lucent space between the separating membranes. Vesicles have formed between the separated membranes. **D.** Three serial sections through a modified connexosome with an inner membrane labeled with Cx43 while the outer membrane has no Cx43 label. All scale bars are 250 nm. Related Video 3 shows the full structures of vesicles in A-D.

The inner and outer membranes have become more separated in 7 of the 56 modified connexosomes and small vesicles were present in this enlarged space (Fig. 3C). The small vesicles were generally not labeled with Cx43, and the diameters averaged 72 nm. This corresponds closely to the reported diameters of intraluminal vesicles (50-80 nm) found in multivesicular endosomes (Murk et al., 2003b; Hanson and Cashikar, 2012; Scott et al., 2014). The inner membrane is identifiable and labeled while the outer membrane is much less labeled (Fig. 3B-D, Video 3). There was a striking example in which the inner and outer membranes were completely separated, and no Cx43 label was present in the outer membrane (Fig. 3D). If the connexons had merely become undocked, the amount of label in the outer membrane should be comparable to the amount in the inner membrane. Therefore, this is evidence for connexon degradation in the outer membrane.

In the remaining 41 of the 56 modified connexosomes, there was an outer membrane similar in diameter to that of unmodified connexosomes that had little to no Cx43 labeling. Also, instead of a single inner membrane labeled with Cx43, there were various different sized vesicles labeled with Cx43 (Fig. 4A and B, Video 3).

**Figure 4.**
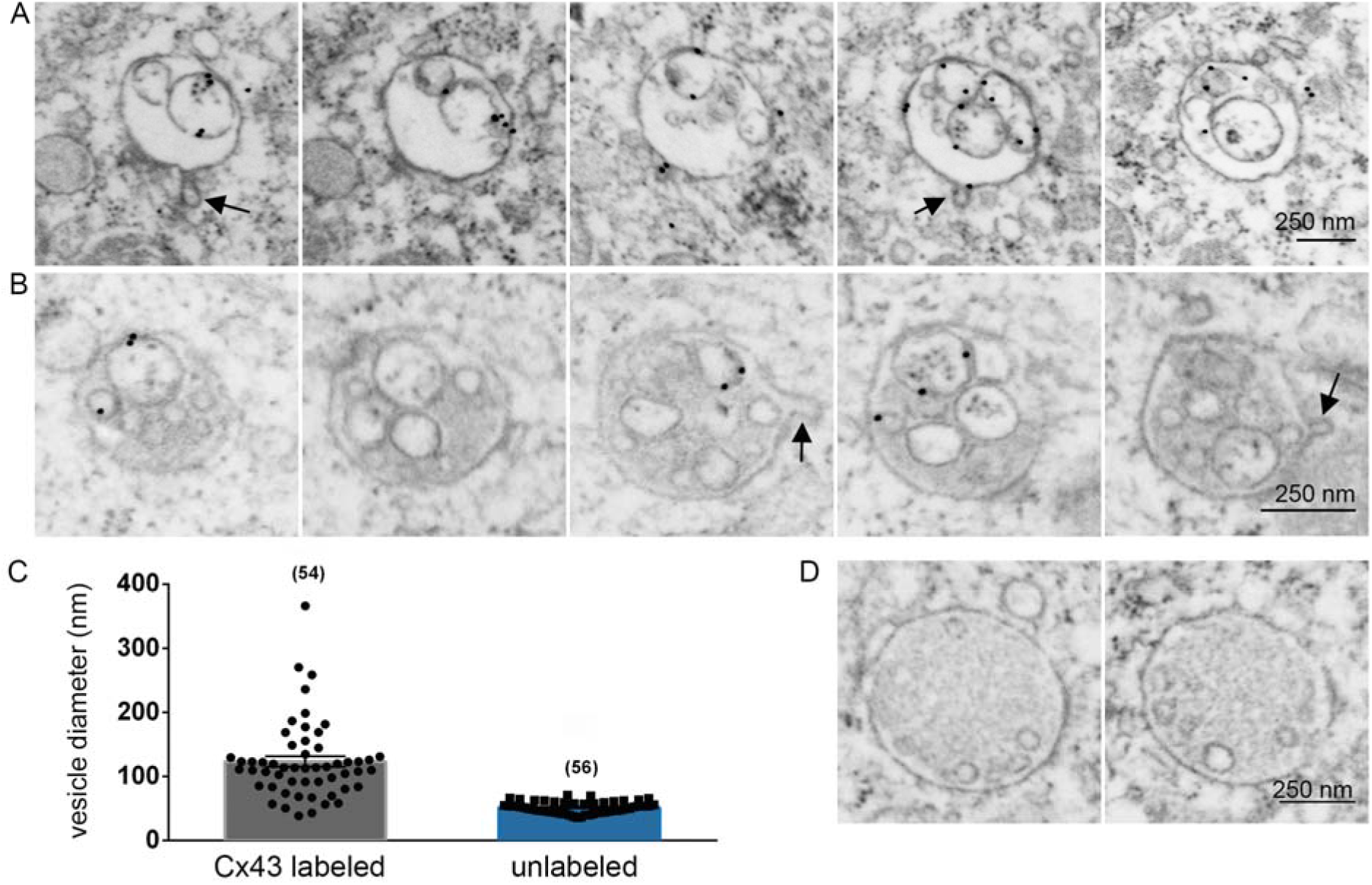
Subdivision of the modified connexosome inner membrane. **A.** Five serial sections through a modified connexosome containing several internal vesicles labeled with Cx43. **B.** Five serial sections through a different modified connexosome with internal vesicles labeled with Cx43. This vesicle has a darker interior than the vesicle in A. In A. and B., arrows indicate short tubules extending from the outer membranes, as is often seen in endosomes. Related Video 3 shows all sections through structures in A and B. **C.** Diameters of Cx43-labeled internal vesicles are more variable in size and larger than unlabeled internal vesicles. 54 internal vesicles labeled with Cx43 were measured within nine modified connexosomes, and 56 unlabeled internal vesicles were measured within nine structures that had no Cx43 labeling. **D.** Two sections through a vesicle with unlabeled intraluminal vesicles, and no Cx43 labeling, as measured for panel C. Scale bars in A,B, and D are 250 nm.

The diameters of the Cx43 labeled vesicles in modified connexosomes averaged 123 nm, and ranged from 38 nm to 366 nm (Fig. 4C). Based on the range of diameters and the presence of Cx43, these seem likely to have formed by fission or outward budding (akin to cytokinesis) of the inner connexosome membrane. For comparison, we also measured small vesicles within structures that lacked Cx43 labeling (Fig. 4D). Unlabeled vesicles were smaller, averaging 56 nm in diameter, with a range of 36 to 70 nm (Fig. 4C).

In addition to subdivision of the inner membrane, some of the modified connexosomes had outer membranes with short tubules extending into the cytosol, (see arrows in Fig. 4A and B), as is often seen in endosomes. (Klumperman and Raposo, 2014).

## Discussion

Luteinizing hormone stimulates the internalization of gap junctions in ovarian granulosa cells (Larsen et al., 1987). We high pressure froze follicles 30 minutes after stimulation and then labeled Cx43 by immunogold staining in serial sections (60 nm thick). Every Cx43 containing structure was categorized and counted in a defined volume of ~30,000 μm^3^. This data allows us to make several novel observations and measurements on connexosome formation and modification.

### Connexosome formation

Live cell imaging studies of several cultured mammalian cell lines expressing Cx43-GFP chimeras showed that either entire gap junctions are internalized, followed by fission (Piehl et al., 2007; Bell et al., 2018), or that a small portion of the gap junction center is internalized (Falk et al., 2009).

The 3D data from high pressure frozen tissue allowed us to look for internalization intermediates and also to make the first systematic measurements of gap junction and connexosome areas. Most gap junctions were disk shaped on a flattened piece of plasma membrane (N = 75). There were 11 invaginated gap junctions, and their average areas were larger than those of the flat gap junctions. It seems likely that the largest gap junctions begin to invaginate and are the source of connexosomes in this tissue. The internalization of whole invaginated gap junctions should produce correspondingly large connexosomes, but connexosomes as a group are smaller than gap junctions. This is consistent with fission occurring soon after internalization.

### Connexosome modifications and fate

The simplest connexosome modification was a partial separation of the inner and outer membrane while the rest of the gap junction is intact. The separated outer membrane bulges out, and lacks Cx43 while the inner membrane retains it (Figures 3B and 3C). It seems likely that a vesicle has fused with the connexosome, perhaps at a bare patch left over from the internalization process (Falk et al., 2009, 2014) If this vesicle fusion leads to a lowering of pH, it could cause the connexons of the gap junction to undock (Falk et al., 2014).

Subsequent modifications seem to involve two different processes. One is complete separation of the two membranes with loss of Cx43 in the outer membrane but retention in the inner membrane. The other process is the appearance of numerous smaller compartments within the boundary of the former connexosome. Small Cx43-free vesicles seem likely to derive from outward budding of the outer membrane; they resemble intraluminal vesicles of multivesicular endosomes. Outward budding or fission of the inner membrane appears to produce Cx43 containing compartments that are somewhat larger than the Cx43 free vesicles.

There is evidence from previous studies of other tissues for connexosome or connexin degradation by autophagy. Autophagosomes engulf internalized gap junctions in the equine hoof wall (Leach and Oliphant, 1984), canine ventricular myocardium (Hesketh et al., 2010), HeLa cells and mouse embryo fibroblasts (Lichtenstein et al., 2010; Fong et al., 2012). In mouse liver cells, connexins are degraded by autophagy (Bejarano et al., 2012). However, we did not observe intermediates resembling a phagophore or an autophagosome in our images. Our data comes from 30 minutes after application of luteinizing hormone, which might be too early for the final stages of Cx43 degradation.

Another possibility is that in ovarian granulosa cells, connexosomes become something more related to multivesicular endosomes. This conclusion was made by Leithe et al (2006) in cultured rat liver epithelial cells that were treated with phorbol ester. Their evidence was based on immunolocalization of endosomal markers and Cx43 by fluorescence and by gold labeling of cryosectioned sucrose-embedded tissue. Falk et al. (2012) later noted that phorbol ester induces phosphorylation of connexin, while in other situations, phosphorylation does not seem to be associated with connexosome formation. Falk et al. (2012) proposed that unphosphorylated connexins target a connexosome for autophagy while phosphorylated connexins lead to multivesicular endosomes. Our observations are consistent with this idea because Cx43 is highly phosphorylated after LH treatment (Norris et al., 2008).

If we had not immunogold labeled sections with Cx43, a structure as seen in Figs. 4A or 4B would likely be identified as a multivesicular endosome. This suggests that in other tissues, some apparent multivesicular endosomes could be modified connexosomes.

### Model for connexosome modification

We propose a sequence of connexosome processing events (Figure 5). The initial event is fusion with a vesicle (Fig. 5, step 1). The vesicle fusion adds unlabeled membrane to the outer membrane, and triggers the undocking of connexons, perhaps by lowering the pH (Falk et al., 2016). The connexons of the outer membrane are degraded, perhaps because the cytoplasm of the host cell can recognize that they are undocked (Fig. 5, step 2) The uncoupled inner membrane undergoes either fission or outward budding to form various sizes of Cx43 containing vesicles (Fig 5, step 3).

**Figure 5.**
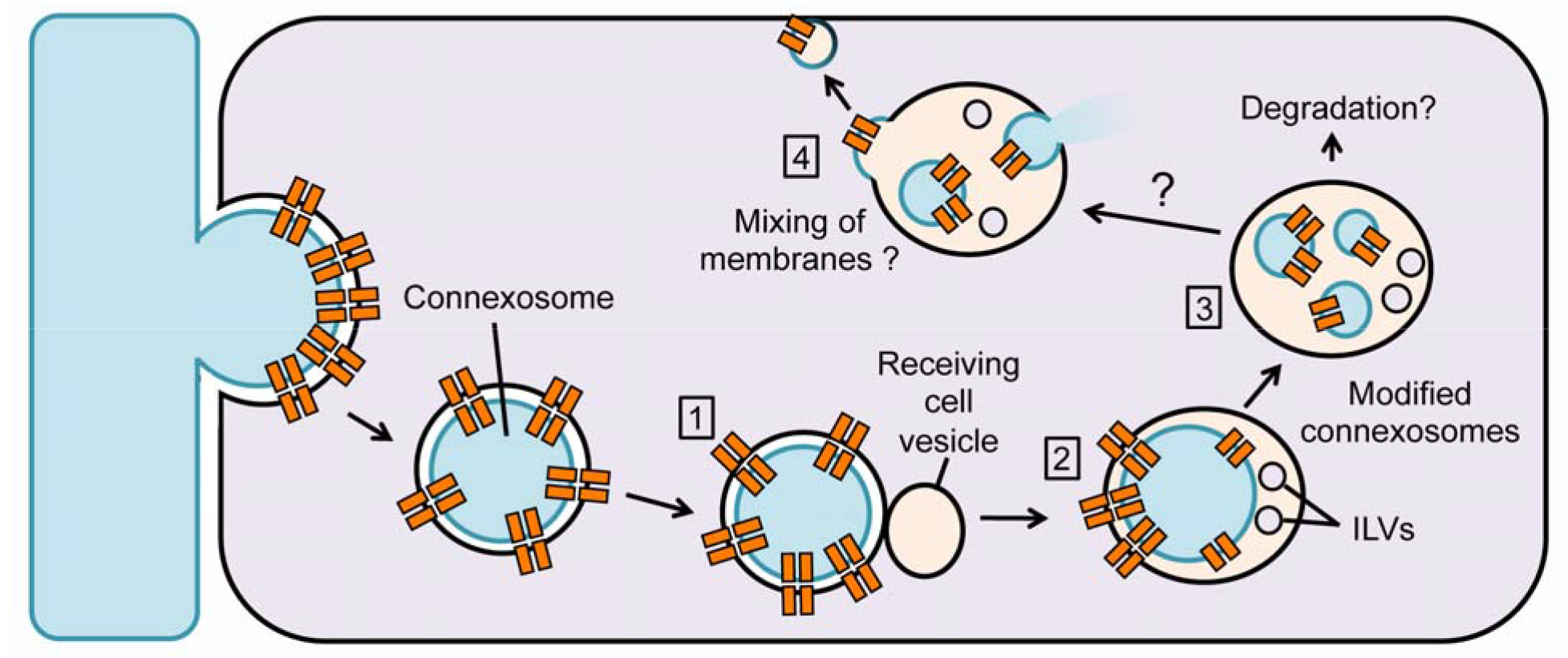
Proposed sequence of connexosome processing. Proposed events after internalization and fission. As in Figure 1, gap junction proteins are represented by orange rectangles. (1). Fusion of a connexosome with a vesicle from the host / recipient cell leads to a local separation of the two membranes (2). The formation of intraluminal vesicles (ILVs) from the outer membrane is also possible. A further modification is the outward budding of the inner membrane that still contains Cx43 (3). Note that Cx43 is decreased or absent in the outer connexosome membrane. While degradation is possible, another possible fate of the modified connexosomes is fusion of the inner vesicles with the limiting membrane of the processed connexosome (4). This would result in release of contents from the other cell and mixing of the plasma membrane from the donor cell with the membrane of the processed connexosome. If some of the donor membrane budded away from the processed connexosome, it could get incorporated into the receiving cell plasma membrane.

Possible fates of the modified connexosomes are degradation by autophagosome formation or direct fusion with a lysosome (Leithe et al., 2006; Falk et al., 2014). However, an alternative possibility is the fusion of the inner vesicles with the modified outer membrane that lacks Cx43 (Fig. 5, step 4). This would result in the mixing of membranes from two cells, and the release of cytoplasm from one cell into another. Evidence for this happening in other cells is discussed in the next section.

What vesicles initially fuse with the connexosomes? Many appear to be clear vesicles, which are consistent with endosomes (Murk et al., 2003a), but a significant fraction of vesicles were dark which is characteristic of lysosomes (Klumperman and Raposo, 2014). There is evidence that lysosomes are not always degradative and can instead function in secretion and signaling (Settembre et al., 2013; Perera and Zoncu, 2016).

While there is solid evidence for clathrin involvement in connexosome formation (Piehl et al., 2007), an ESCRT driven process could also help explain some of our observations. ESCRT machinery for outward budding could become attached and activated on the donor side of the gap junction. There is evidence that ESCRT machinery can bind to ubiquitinated connexins (Auth et al., 2009). If the machinery were transferred within the connexosome, it could generate single membrane vesicles by outward budding or fission of the internal membrane after separation of the inner and outer connexosome membranes.

### Cytoplasmic and membrane transfer by connexosomes

The channel properties of connexins are well established for the exchange of small molecules between cells (Nielsen et al., 2012). We discuss here how connexins, by way of forming connexosomes, may also facilitate cytoplasmic and membrane transfer between cells.

First of all, our serial section images provide conclusive evidence for transfer of mitochondria and other organelles via connexosomes. A transferred mitochondrion could affect physiological processes of the host cell. In a mouse model for acute lung injury, Cx43-dependent mitochondrial transfer from bone marrow derived stroma cells rescued injured lung alveolar epithelial cells (Islam et al., 2012). The authors suggested that mitochondria were transferred in some way by microvesicles; our observations provide a clear Cx43-dependent mechanism for transfer.

Our observations demonstrate only the transfer of an organelle in an enclosed double membrane. If the double membranes were to fuse (or the inner membrane vesicles were to fuse with the outer membrane), this would result in mixing of cytoplasm (e.g. release of mitochondrion in the above example) and membranes (Figure 5, step 4). It is established that the endosome membrane can fuse with a vesicle within it (Bissig and Gruenberg, 2014). This occurs with enveloped viruses such as vesicular stomatitis virus (Le Blanc et al., 2005) or coronavirus (Grove and Marsh, 2011), resulting in the release of nucleocapsids into the cytoplasm. There is now direct evidence that this also occurs with extracellular vesicles (Joshi et al., 2020), resulting in release of cargo to the cytoplasm.

Membrane mixing involving connexosomes might explain a previous finding in immune cells. Macrophages transfer MHC II bound antigens to dendritic cells in the gut to establish oral tolerance, and this depends on the presence of Cx43 (Mazzini et al., 2014). Without specifying the role of Cx43, the authors raise the possibility that transfer occurs via trogocytosis. In trogocytosis, like connexosome formation, a part of the plasma membrane and cytoplasm of one cell is internalized into another (Joly and Hudriser, 2003); a fusion between outer and inner compartments, followed by budding of the combined membranes would be required to achieve antigen transfer and presentation. We suggest that the MHC II-antigen complex is transferred from the macrophage via connexosomes followed by fusion of inner and outer membranes to deliver it to the dendritic cell’s endosomal system. Cell-cell communication by way of connexosome formation and processing may be widespread and warrants further investigation.

## Materials and Methods

### Mice

C57Bl/6J female mice (*Mus musculus*) from The Jackson Laboratories (Bar Harbor, ME) were used in this study. The use of mice was in accordance with guidelines published by the National Institutes of Health and approved by the UConn Health Institutional Animal Care and Use Committee. Mice were housed in a temperature and humidity controlled environment with water and food available ad libitum.

### Antibodies

Anti-Connexin 43 polyclonal antibody produced in rabbit (#C6219, Sigma-Aldrich, St. Louis, MO) was used at 1:100 for a final concentration of ~7 micrograms/ml. Secondary antibody, 10 nm gold particle conjugated goat anti-rabbit IgG F(ab’) 2 fragment, (#25362, Electron Microscopy Sciences, Hatfield, PA) was used at a dilution of 1:20.

### Preparation of follicles for post-embedding immunogold labeling

Ovarian follicles, 360-400 micrometers in diameter, were manually isolated from 25 day-old mice. The follicles were cultured on Millicell membranes (Millipore, PICMORG50) in MEM alpha medium as described by Norris et al., 2008, with the replacement of 3 mg/ml bovine serum albumin (MP Biomedicals, #103 700) for serum.

After 25 hours of culture, follicles were treated with 10μg/ml ovine luteinizing hormone (oLH-26, National Hormone and Pituitary Program) for 30 minutes, then transferred to brass specimen carriers and high pressure frozen with an EMPACT 2 (Leica Biosystems, Buffalo Grove, IL).

From this point, we followed the procedure of Rubio and Wenthold (1997), with some modifications. Samples were freeze-substituted with 2.5% uranyl acetate (Electron Microscopy Sciences) in dry methanol for 32 hours at −90° C in an AFS 2 freeze substitution unit (Leica Biosystems). The temperature was then raised 5° C per hour to −45° C. Samples were then rinsed in methanol, and infiltrated with Monostep Lowicryl HM-20 (Electron Microscopy Sciences, Hatfield, PA) in increasing concentrations of 1:1, 2:1, and pure Lowicryl HM20 for two hours each. After an overnight infiltration with Lowicryl HM-20, samples were polymerized in a fresh change of Lowicryl under ultraviolet light for 33 hours at −45° C. Polymerization continued under UV light as the temperature in the AFS 2 was increased by 5° per hour to 0° C, then held at 0° C for 33 hours more. When samples were removed from the AFS 2, they were pink in color and left to polymerize at room temperature until the pink hue was gone two days later.

### Collection of serial sections

Ultrathin sections (60 nm) of Lowicryl HM20 embedded follicles were cut on a UC-7 ultramicrotome (Leica Biosystems) with a diamond knife (Diatome, Hatfield, PA). The sections were picked up by an automated tape collector on glow-discharged kapton tape (Terasaki et al., 2013; Kasthuri et al., 2015; Baena et al., 2019).

### Immunogold staining of serial sections

For immunostaining, ribbons of follicle sections on kapton tape were cut to lengths of approximately three inches and attached to a sheet of parafilm with double-sided carbon tape (Electron microscopy Sciences #77816).

Sections were rehydrated with 1X PBS (Life Technologies, Grand Island, NY) and blocked in 5% normal goat serum (Invitrogen, Frederick, MD) in a solution of 1% bovine serum albumin in PBS.

Following an overnight incubation at 4°C in primary antibody, sections were rinsed three times for 5 minutes each in PBS, then rinsed once in 1% BSA in PBS. Next, secondary antibody diluted at 1:20 was applied to sections for one hour at room temperature. Sections were then rinsed with 1X PBS followed by Milli-Q filtered water and dried overnight. Sections were placed back in their original order and post-stained with 5% uranyl acetate in 50:50 methanol: water for 7 minutes, then rinsed generously in water.

### Imaging serial sections of tissue with scanning EM

Immuno-labeled sections on tape were attached to a 10 cm diameter silicon wafer (University Wafer, South Boston, MA) with double-sided carbon adhesive tape (Electron Microscopy Sciences). Wafers were carbon coated (Denton, Moorestown, NJ) and first imaged on a Sigma field emission scanning electron microscope (Zeiss, Thornwood, NY) using a backscatter detector as described in Norris et al., 2017. Two volumes of mural granulosa cells were imaged at 5 nm/pixel resolution, with a field of view of 60 square micrometers.

### Analyzing image stacks

Original low-resolution images obtained on the Zeiss Sigma were aligned with the Register Virtual Stack Slices macro (FIJI), then larger files were aligned and diced for convenient viewing with a custom program (Piet, provided by Duncan Mak and Jeff Lichtman, Harvard University). These files were used to track all Cx43-labeled internalized structures. When internalized structures looked complex, they were re-imaged using a higher resolution electron microscope as described below.

### High-resolution images on FEI Verios 460L

Higher resolution images of the sections were taken on an FEI Verios 460L field emission scanning microscope with a backscatter detector. Using immersion mode, sections were imaged at 5.0 kV, with a beam current of 0.4 or 0.8 nA. Images had a dwell time of 3.0 microsec, with a line integration of 5 or 10, and drift correction on. Image sizes of 2048 x 1768 were taken with horizontal field widths between 2.76 to 7.89 micrometers to generate images with resolutions between 1and 4 nm/pixel. Under these conditions, we could detect the characteristic pentalaminar appearance of gap junctions in some sections. To minimize beam damage, it was beneficial to first use a ‘beam shower’ over the sections by scanning at a lower magnification, with a higher beam current, and a faster scan rate for 10-20 seconds.

### Measuring surface areas

Image stacks were created by aligning serial images with the ‘Register Virtual Stack Slices’ macro in Fiji. Stacks were imported into the TrakEM2 program in Fiji software to measure surface areas. Separate ‘area lists’ were created for reconstructions and measurements of individual gap junction plaques or connexosomes. Images of 15nm/pixel were traced. Using the ‘measure’ tool, the column labeled ‘AVG-s’ (Average smooth) was used as an estimate of surface area for both connexosomes and gap junction plaques. Default units of pixels squared were converted into micrometers squared. To calculate the surface areas of gap junction plaques, the ‘AVG-s’ values were first divided by two, as the initial value represented both faces of the gap junction plaque.

## Supplementary Material

Three videos are included as supporting material. All videos show serial sections through ovarian granulosa cells immunogold-labeled for Cx43. Video 1 shows several connexosomes and other vesicles labeled with Cx43. Video 2 shows an invaginated gap junction allowing organelles from one cell to protrude into another and the full structures of connexosomes containing cytosol and organelles. Video 3 shows various connexosome modifications.

## Acknowledgements

We thank Rindy Jaffe and Matthias Falk for critical review of the manuscript and useful discussions. We thank Art Hand, Maya Yankova, and Maria Rubio for technical advice. We thank Valentina Baena, Tracy Uliasz, and other members of the Jaffe and Terasaki labs for technical assistance and help with collecting samples. This work was funded by the Fund for Science. The authors declare no conflicts of interest.

## Author Contributions

Rachael Norris contributed to conceptualization; data collection, curation and analysis; visualization, and writing and editing of the original draft. Mark Terasaki contributed to conceptualization, supervision of the work, funding acquisition, software and resources for data collection; and writing and editing of the original draft.

## Video Legends

**Video 1. Multiple structures labeled with Cx43 polyclonal antibody.** Serial sections through an ovarian granulosa cells showing a variety of double membrane vesicles labeled with an antibody to Cx43 and 10nm gold. Other types of vesicles are also labeled with Cx43 (yellow arrows). The scale bar is 250nm.

**Video 2. Organelle transfer by gap junction internalization.** Serial sections through ovarian granulosa cells labeled with anti-Cx43 and 10nm gold. Corresponding to Figure 2A, the cytoplasm of a cell protruding into another cell is shaded in blue. There is a multivesicular endosome and a mitochondrion protruding into the cell as well. Scale bar is 500nm, as in Figure 2A. Corresponding to Figure 2B, a connexosome contains a multivesicular endosome and a mitochondrion. Corresponding to Figure 2C, another connexosome contains smaller vesicles and other membranes. Scale bars are 250nm, as in Figures 2C and 2C.

**Video 3.** Connexosome modifications of the outer and inner membranes. Serial sections through ovarian granulosa cells labeled with anti-Cx43 and 10nm gold. Corresponding to Figures 3 and 4, the full structures of the modified connexosomes in Figure 3A-D and in Figures 4A and B are shown. Scale bars are 250nm.

## Notes

### Competing Interest Statement

The authors have declared no competing interest.

